# Control of spontaneous activity patterns by inhibitory signaling in the developing visual cortex

**DOI:** 10.1101/2020.02.21.959262

**Authors:** Alexandra H. Leighton, Gerrit J. Houwen, Juliette E. Cheyne, Paloma P. Maldonado, Fred De Winter, Christian Lohmann

**Author notes:** Author contributions, Conceptualization: A.H.L., J.E.C. and C.L.; Methodology: A.H.L., J.E.C. and G.J.H.; Formal Analysis: A.H.L., P.P.M and J.E.C.; Resources: F. DW.; Investigation: A.H.L., J.E.C., P.P.M. and G.J.H.; Writing: A.H.L. and C.L.

## Abstract

During early development, even before the senses are active, bursts of activity travel across the nervous system. This spontaneously generated activity drives the refinement of synaptic connections, preparing young networks for patterned sensory input. Synaptic fine-tuning relies not only on the presence of spontaneous activity, but also on the specific characteristics of these activity patterns, such as their frequency, amplitude and synchronicity. Here, we provide evidence that these crucial characteristics are shaped by the relative balance of excitation and inhibition, where patterns with distinct characteristics have different excitatory/inhibitory ratios. Inhibition can control whether cells participate during a spontaneous event, as pharmacogenetic suppression of the somatostatin (SST) expressing subtype of inhibitory interneurons increased cell recruitment and lateral spread of events.

## Introduction

In order to survive, very young animals must be able to interact with their environment as soon as they start receiving useful information about their surroundings. To prepare for this, the nervous system is thoroughly organized even before the onset of sensory input (for instance, when the eyes or ears open in mice). Young cells are initially shaped into rough networks by molecular guidance cues, and these connections are subsequently refined by activity dependent processes (Cline, 2003; Sanes and Yamagata, 2009). The necessary activity is largely generated by spontaneously depolarizing cells in the sensory organs and the brain (Blankenship and Feller, 2010), which initiate and propagate patterned ‘training’ activity across the developing network to strengthen well-targeted synapses and weaken others.

The best studied example of refinement through spontaneously generated activity is in the mouse visual system, where the neonatal retina generates bursts of activity which travel downstream and refine retinotopic maps as well as the segregation of contra- and ipsilateral afferents through Hebbian and non-Hebbian mechanisms (Kirkby et al., 2013). As a result of this fine-tuning, neurons in the mouse visual cortex can respond with striking acuity to visual information as soon as the eyes open at postnatal day (P)14 (Cang et al., 2005; Ko et al., 2013; Rochefort et al., 2011; Zhang et al., 2012).

Different patterns of spontaneous activity occur throughout the central nervous system. Such patterns of activity can be described and distinguished by their characteristics: the frequency with which activations occur, synchronicity of cell firing, number of action potentials fired, and number of cells that participate in each event (Ackman and Crair, 2014; Allene and Cossart, 2010; Colonnese and Phillips, 2018; Kerschensteiner, 2014; Luhmann and Khazipov, 2018).

A growing body of work has shown that merely the presence of activity is not sufficient for refinement, but that these specific characteristics encode and transmit essential information required by the brain to develop normally (Kirkby et al., 2013; Leighton and Lohmann, 2016). For instance, if retinal waves are too large, retinotopic map refinement is prevented, whereas eye-specific segregation can be impaired by changes in event frequency (Burbridge et al., 2014; Xu et al., 2011). In the primary visual cortex (V1), two types of activity are reported - one type driven by retinal waves (Ackman et al., 2012; Gribizis et al., 2019) and reduced upon enucleation, and activity which occurs independently of retinal activity (Gribizis et al., 2019; Hanganu et al., 2006; Siegel et al., 2012). These two types of events have different characteristics that can be readily quantified with two-photon calcium imaging. Retinally-driven events activate relatively few neurons and are therefore referred to as low participation events (‘L-events’). Their retinal origin, combined with their comparative sparsity of activation, gives them the potential to shape the network according to the organization of the eye. In contrast, those events which occur independently of retinal manipulations cause high-amplitude activity in almost all cells in a large field of view. These high participation events (‘H-events’) may allow neurons to perform synaptic homeostasis, bringing synaptic strengths back to a workable range (Siegel et al., 2012).

How does the immature brain control the characteristics of these activity patterns? In adults, we have an increasing understanding and appreciation of the strong regulation of network activity by inhibitory interneurons, as well as the specific roles of interneuron subtypes (Kepecs and Fishell, 2014; Markram et al., 2004; Tremblay et al., 2016; van Versendaal and Levelt, 2016). In neonatal animals however, our understanding of GABAergic control is less thorough. During very early development, GABA is thought to act as a depolarizing neurotransmitter, switching to its adult inhibitory role as cells mature (Cherubini et al., 1991). However, from the end of the first postnatal week GABAergic cells exert an inhibitory effect on the cortical network (Che et al., 2018; Kirmse et al., 2015; Minlebaev et al., 2007; Valeeva et al., 2016) and are therefore prime candidates to control spontaneous event patterning even before eye opening. Furthermore, GABAergic signaling is required for activity-dependent wiring of the developing cortex (Che et al., 2018; Duan et al., 2019; Marques-Smith et al., 2016; Modol et al., 2019; Oh et al., 2016; Tuncdemir et al., 2016). Here, we investigated how inhibitory signaling shapes spontaneous event patterning in V1 before eye opening. We confirm that L- and H-events have very different characteristics. We find that L- and H-events have different excitation/inhibition dynamics, where stronger excitation underlies H-event properties and L-events are under tighter inhibitory control. By manipulating inhibitory signaling, we show that specific characteristics of spontaneous activity can be changed. Together, our results suggest that those characteristics of spontaneous activity that give them the power to fine-tune the developing network, such as frequency, amplitude and specificity, are controlled by the balance of excitation and inhibition during the second postnatal week.

## Results

### L and H events are two distinct network activity patterns at both cellular and population level

We previously described L- and H-events using two-photon calcium imaging (Siegel et al., 2012). To understand how L- and H-characteristics match up to other reports of spontaneous activity, we measured their properties at a range of different scales. First, we observed how L- and H-events differ at single-cell level by recording action potential firing from primary visual cortex (V1) layer 2/3 neurons using whole cell recordings *in vivo* in mouse pups between P8 and P12. We used simultaneous two-photon calcium imaging to identify L- and H-events (Fig 1A, B). Neurons fired significantly more action potentials during H-events than during L-events, in agreement with their higher calcium transient amplitude (Fig 1C). We detected no significant difference between the inter-spike intervals within a burst (Supplementary Fig 1A). Instead, the duration of spiking in H-events was significantly longer than in L-events (Supplementary Fig 1B). L-event duration (mean 4.6 seconds) was similar to the duration of retinal waves (summarised in Torborg and Feller, 2005), in line with their largely retinal origin.

To explore the separation of these events into two distinct groups (L and H) in a manner independent of calcium imaging, we performed hierarchical clustering using the number of action potentials fired and the duration of bursts. Silhouette analysis revealed an optimum of two clusters (Supplementary Fig 1C). This split (Supplementary Fig 1D) corresponded well (76%
overlap) to our original definition of L- and H-events, where events with between 20 and 80% participation classified as L-events, and those over 80% network participation were identified as H-events (Siegel et al., 2012). We therefore continued to use participation to distinguish Land H-events during two-photon calcium imaging.

**Figure 1.**
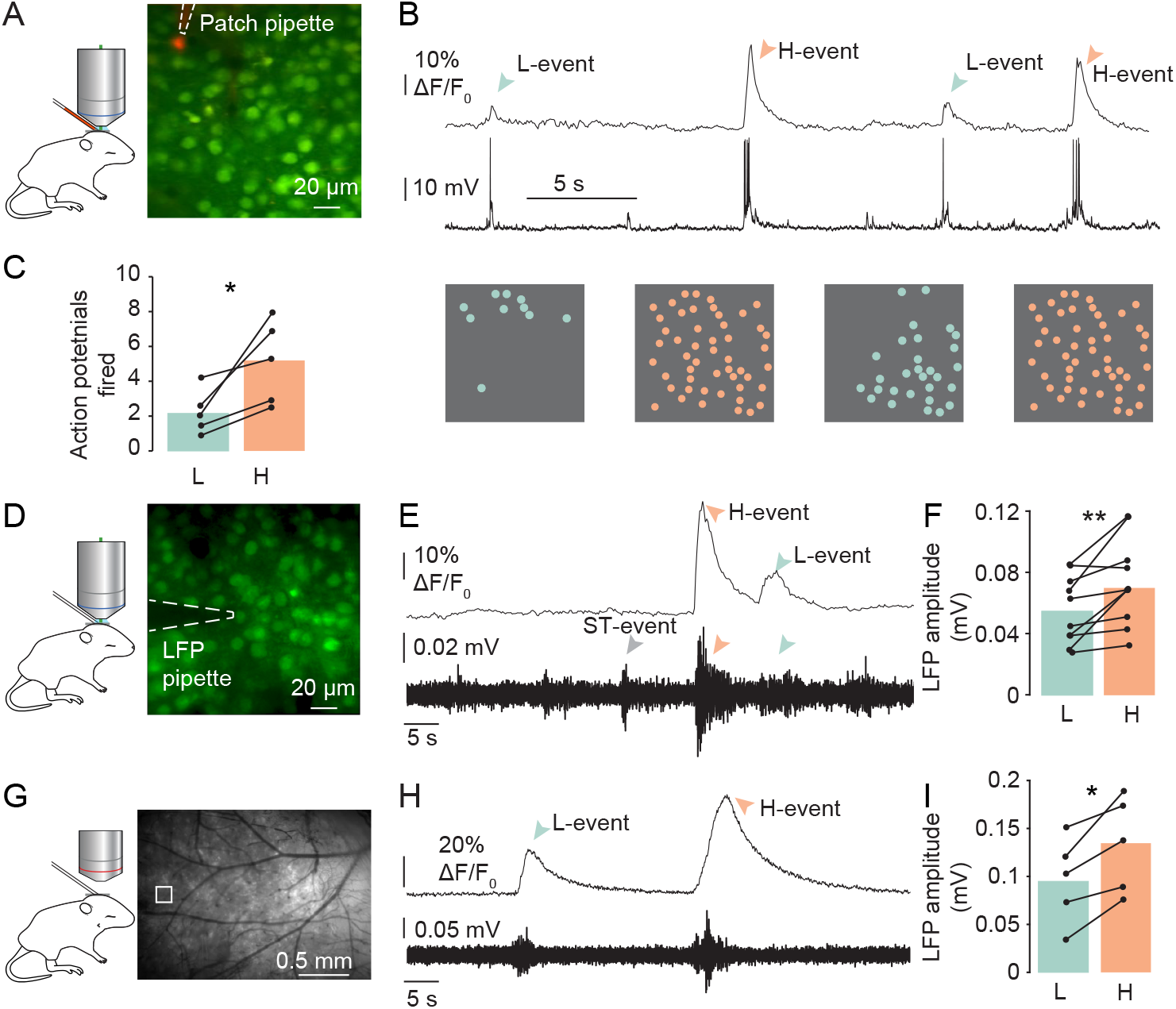
Distinct spontaneous activity patterns occur in the visual cortex during the second postnatal week A. Cells in V1 L2/3 were loaded with the calcium indicator Oregon Green Bapta 1-AM (green) to track spontaneous activity patterns. The recorded cell was filled with Alexa 594 via the patch pipette for identification (red). B. L-events (blue arrowheads) and H-events (orange arrowheads) shown as a calcium trace (average activity of all imaged cells) with simultaneous whole-cell current-clamp recording. The cells that participated in each event are displayed as filled circles below. C. More action potentials were fired during H-events than during L-events (p = 0.03, paired t-test, n = 5 animals). D. Cells in V1 L2/3 loaded with the calcium indicator Oregon Green BAPTA 1-AM (green). The shadow of the LFP pipette used is visible. E. Averaged calcium trace of imaged cells (above) with simultaneous LFP recording (below). The LFP reveals subthreshold (ST) events that are not visible in the calcium imaging (grey arrowhead). F. The LFP amplitude was significantly higher during H-events than during L-events (p = 0.009, paired t-test, n = 10 animals). G. Epifluorescence imaging of the visual cortex using GCaMP-6s was used to image a larger field of view during LFP recordings. Small white square indicates field of view in two-photon. H. Mean calcium responses during L- and H-events (above) and simultaneous LFP recording (below). I. The LFP amplitude was significantly different between L- and H-events defined in wide-field imaging (p = 0.014, paired t-test, n = 5 animals).

We then proceeded to population-level measurements, using local field potential (LFP) recordings. Spindle bursts are a commonly described pattern of spontaneous activity in the LFP in visual (Hanganu et al., 2006) and somatosensory (Khazipov et al., 2004) cortex, characterized by low frequency spindle-shaped oscillations. To link these findings to our description of network events, we recorded the LFP during two-photon calcium imaging of L- and H-events. We observed spindle bursts in the LFP during both L- and H-events (Fig 1D, E). The LFP also revealed ‘subthreshold’ events that were not associated with calcium transients within the field of view (0.026 mm^2^), presumably the result of events occurring outside the field of view of two-photon imaging. LFP amplitude was significantly higher during H-events than during L-events (Fig 1F), in line with the greater synchronicity of H-events reported previously (Siegel et al., 2012). The peak frequency component did not differ between L- and H-events (Supplementary Fig 1E), consistent with the similarity of action potential inter-spike-intervals in both event types.

Accurate refinement of projections requires activity with limited spatial spread (Burbridge et al., 2014; Xu et al., 2015). As H-events activate almost all neurons in the two-photon field of view, we used wide field imaging to record a much larger part of the visual cortex (4.9 mm^2^, Fig 1G, H), allowing us to quantify the spatial spread of events. To understand whether we could observe L- and H-events on this scale, we performed hierarchical clustering on the wide-field events. As L- and H-events, when measured with two-photon microscopy, are distinguished by their amplitudes and participation, we used mean calcium amplitude and event size as wide-field proxies of these parameters. Silhouette analysis again revealed an optimal separation at two clusters (Supplementary Fig 1F), splitting the data into events occurring locally and with low amplitudes and events with large spatial spread (Supplementary Fig 1G) and with high amplitudes. The LFP corroborated this difference, as H-events had a significantly higher LFP amplitude than L-events (Fig 1I). L-events also occurred at a much higher frequency (Supplementary Fig 1H). To compare L- and H-type activity between wide-field and two-photon microscopy, we measured event frequency at an area corresponding to the size of the field of view of our two-photon microscope. We found that H-events occurred at a frequency of 0.6 ± 0.25 events per minute, similar to the previously determined frequency in two-photon experiments (0.5 per minute; Siegel et al., 2012). For L-events, we saw an average of 0.37 ± 0.12 events per minute.

Taken together, these results confirmed that two types of network activity can be distinguished at both single neuron and population level in the visual cortex during the second postnatal week, and that they differ by the number of spikes fired, their synchronicity (reflected in LFP amplitude) and their lateral spread. These properties may depend on the relationship between excitatory and inhibitory signaling during network activity, as the number of spikes fired by a given neuron and the number of recruited neurons increase when the excitatory drive is high or inhibition is low. Furthermore, reducing inhibition increases network synchronization (Valeeva et al., 2010) whereas fast, correlated inhibition and excitation can reduce synchronization (Harris and Thiele, 2011). Finally, inhibition can reduce the number of neurons recruited to an event, limiting activity spread (Stefanelli et al., 2016). We therefore hypothesized that L- and H-events have different excitation/inhibition dynamics, where stronger excitation underlies H-event properties and L-events are under tighter inhibitory control.

### The excitation/inhibition ratio is higher during H-events than during L-events

To compare excitatory and inhibitory output during spontaneous activity, we used mice in which neurons expressing the GABA production enzyme Glutamate Decarboxylase 2 (GAD2) were labelled with the red fluorescent protein tdTomato (Fig 2A). We found that GAD2^+^ cells participated in both L- and H-events. 16.5 ± 5.9% of active neurons in an event were GAD2^+^ (Fig 2B), corresponding to the average percentage of imaged neurons that expressed GAD2 (17.1 ± 5.9% across all ages).

During L-events, the mean amplitudes of calcium responses in GAD2^+^ and GAD2^-^ cells were very similar (Fig 2C, E). During H-events, GAD2^+^ cells showed slightly, but significantly, lower amplitudes than GAD2^-^ cells (Fig 2D, F). Consequently, the ratio between H- and L-event amplitude was significantly lower in GAD2^+^ cells (Fig 2G). Using data from the simultaneous network imaging and patch clamp experiments in Fig 1A, we fitted a linear function to the relationship between the average cell amplitude in an event and the number of action potentials fired by the neuron (Supplementary Fig 1I). This translated the calcium transient into a difference of one action potential between GAD2^+^ and GAD2^-^ output. These data suggested that both GAD2^+^ and GAD2^-^ cells fired more action potentials during H-events than during L-events, but that during H-events, GAD2^+^ cells fired fewer action potentials than GAD2^-^cells, though this difference was relatively small.

**Figure 2.**
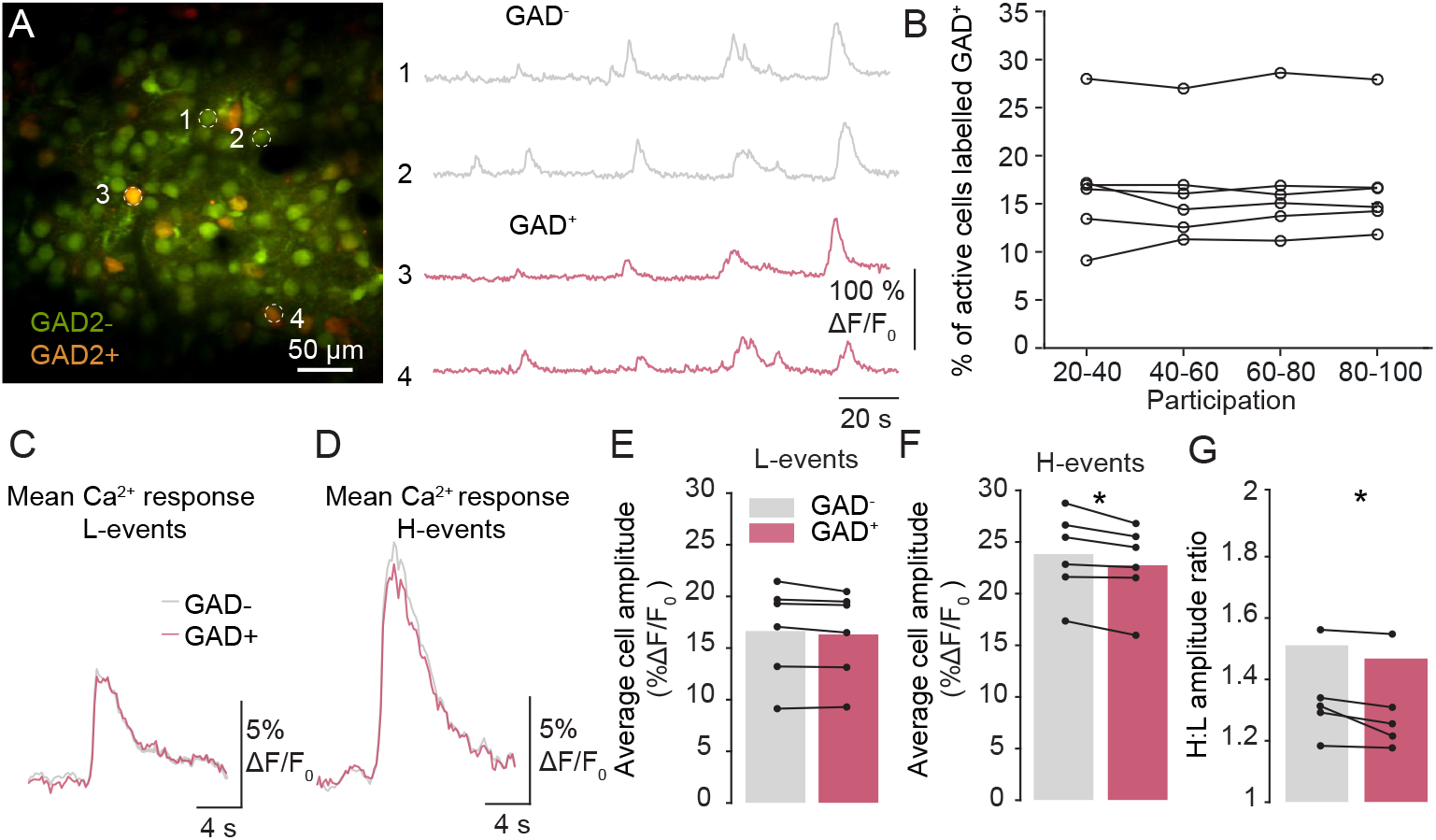
Excitatory/inhibitory output balance is different during two distinct types of spontaneous network events A. Left: Cells in V1 L2/3 of a GAD2-cre x CAG-tdTomato mouse. All cells were loaded with the calcium indicator Oregon Green BAPTA 1-AM (green). GAD+ cells expressed tdTomato (red). Right: four traces from example GAD-(above) and GAD+ (below) cells. B. The percentage of participating cells that expressed GAD remained constant over all participation bins for each animal (n = 6). C. The mean calcium response of all active cells during L-events recorded in an example animal. D. The mean calcium response of all active cells during H-events recorded in an example animal. E. In L-events, the average cell amplitude does not differ between cells that do and do not express GAD (n.s., paired t-test, n = 6 animals). F. In H-events, GAD+ cells have a significantly lower amplitude than GAD-cells (*p < 0.05, paired t-test, n = 6 animals). G. The ratio between H:L event amplitude is slightly but consistently lower for GAD+ than GAD-cells.

Next, we examined whether the excitatory/inhibitory balance during L- and H-events differed on the level of synaptic inputs received by each neuron. We performed *in vivo* whole-cell recordings in voltage-clamp mode to measure excitatory and inhibitory synaptic input currents onto neurons (Figure 3). Simultaneous calcium imaging of the network allowed identification of L- and H-events. As reported previously, events involved both glutamatergic and GABAergic synapses (Fig 3A, Colonnese, 2014; Hanganu et al., 2006). In line with the higher network participation in H-events, both the excitatory and inhibitory charge transferred were larger during H-events than during L-events (Fig 3B-D). The ratio of charge transferred between Hand L-events was significantly larger for excitatory currents than for inhibitory currents (Fig 3E), demonstrating a difference in the amount of excitation relative to inhibition during network events.

Next, we trained a random forest classifier (Breiman, 2001) using only electrophysiological measurements to decode whether an event was an L- or an H-event, using the peak current amplitude, mean current amplitude, total charge transferred and duration of either inhibitory or excitatory current inputs. Training event classification was based on the original definition of 20-80% (L-events) and over 80% participation (H-events) in network events obtained with two-photon calcium imaging. For all animals, the classifier trained on the excitatory data performed significantly better than the classifier trained on the inhibitory data, as demonstrated by the larger area under the curve of the receiver-operator characteristics (ROC) curve (Fig 3F, G).

These results demonstrated that during H-events, neurons not only received more input overall but specifically received a higher ratio of excitatory to inhibitory inputs when compared to L-events.

**Figure 3.**
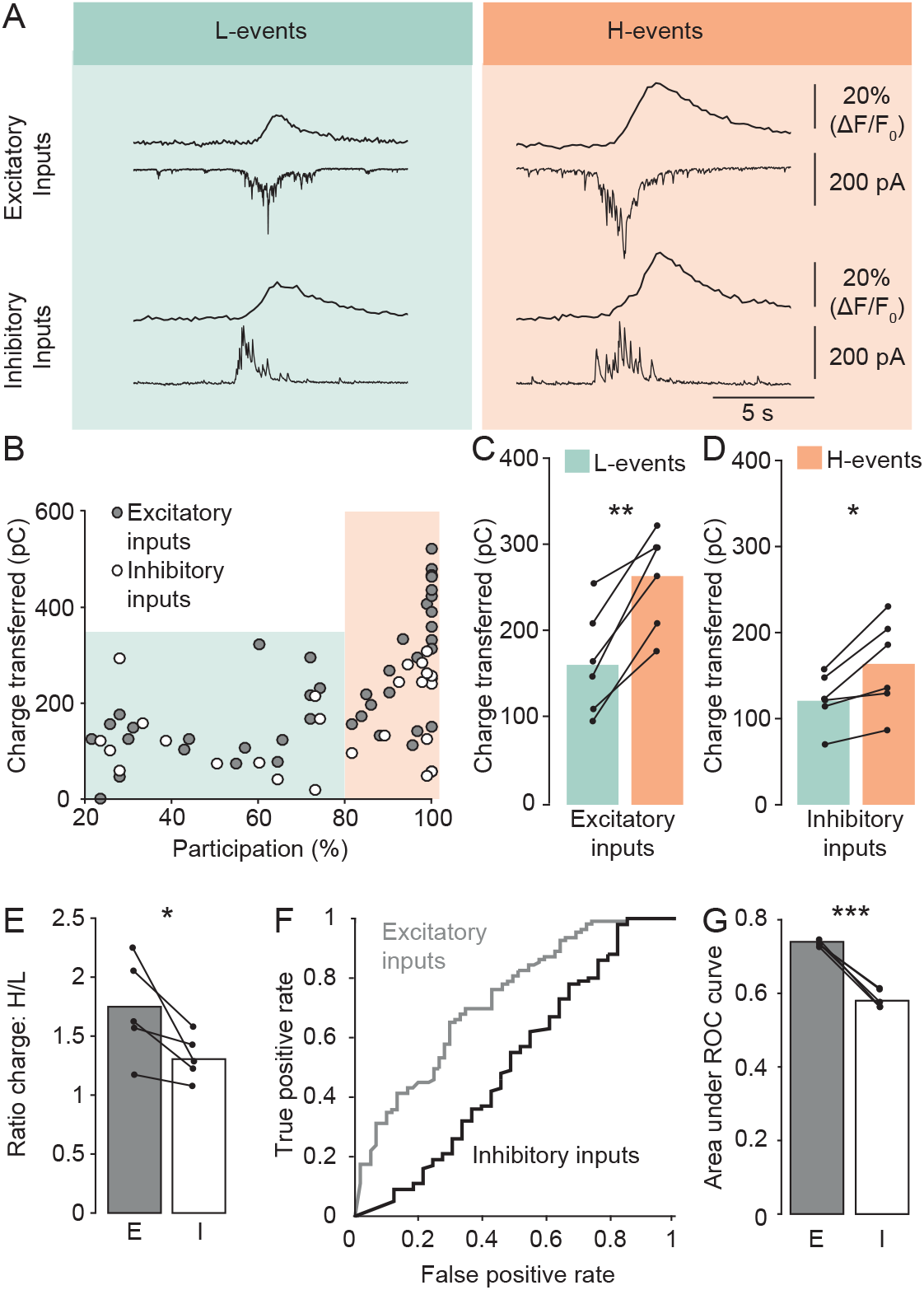
Excitatory inputs contain more information about the type of network activity than inhibitory inputs A. In vivo voltage-clamp recordings combined with network calcium imaging in layer 2/3 V1 neurons. Excitatory and inhibitory inputs were measured alternatingly by switching the holding potential to the reversal potential of inhibitory or excitatory currents, respectively. Four events in an example cell, showing inhibitory and excitatory inputs received by the cell during an L-event and an H-event. Average calcium traces are shown from all imaged cells (above). Current traces (below) represent synaptic inputs onto the recorded neuron. B. Absolute transferred charges of excitatory and inhibitory input currents during network events of different participations in one cell. C. The excitatory charge transferred was significantly higher during H-events than during L-events (p = 0.001; paired t-test, n = 6 animals). D. The inhibitory charge transferred was significantly higher during H-events than during L-events (p = 0.016, paired t-test, n = 6 animals). E. The H:L ratio of charge transferred was significantly larger for excitatory than for inhibitory inputs (p = 0.036, paired t-test, n = 6 animals). F. Example receiver operating characteristic (ROC) curve for one animal, showing a random forest classifier trained on excitatory or inhibitory data. The larger area under the curve for excitatory inputs indicates a higher success rate of classifying L- and H-events correctly when excitatory data was used compared to inhibitory data. G. Quantification of the area under the ROC curve for all animals. The area under the curve is significantly higher for classifiers trained on excitatory input data than for those trained on inhibitory input data (p = 0.0002, paired t-test, n = 5 animals).

### Excitatory and inhibitory signaling are essential for patterned spontaneous activity

The above experiments confirmed that activity patterns with distinct properties and origins had different excitatory/inhibitory balances. From the synaptic input data, the difference between L- and H-events seemed to arise from an increase in excitation, rather than a change in the amount of inhibition. To test the importance of inhibitory signaling at this age, and to establish the causality of the relationship between excitatory/inhibitory balance and event properties, we tested whether changing the excitatory/inhibitory balance affected activity patterns using pharmacological manipulations and two-photon calcium imaging (Figure 4). Applying the AMPA and NMDA receptor antagonists NBQX (50 μM) and APV (100 μM) locally on the cortical surface abolished spontaneous activity (Fig 4A, C). Blocking GABA_A_ receptors with picrotoxin (PTX, 200 μM) greatly increased the frequency of spontaneous activity, confirming the inhibitory effect of GABAergic signaling on the network (Fig 4B, D). Consistent with reduced inhibition, the peak calcium transient amplitude increased after PTX (Fig 4E). Finally, PTX application significantly increased the participation of each event, implicating inhibitory signaling in control of neuron recruitment (Fig 4F).

These results indicated that glutamatergic signaling drives spontaneous activity whereas GABAergic signaling shapes its properties in V1 during the second postnatal week. Reducing inhibition increased the frequency and caused network activity properties to switch to H-like characteristics, supporting the idea that the characteristics of H-events are generated by the excitatory/inhibitory ratio being tipped towards excitation.

### Somatostatin cell suppression reduce event specificity

Our results so far showed that during the second postnatal week, the network can maintain different amounts of inhibitory signaling relative to excitation, and that this balance and its endogenous modulation determine specific activity pattern characteristics.

The retinally-driven L-events are characterized by their low participation, activating subsets of neurons in the field of view. This restricted participation of L-events may allow them to mediate refinement of the network. In contrast, H-events are highly synchronized in time and drive substantial areas of the cortex to fire large numbers of action potentials, and presumably lack the specific information required for local refinement.

Given that L-events arise from balanced inhibition with excitation, we aimed to find a manipulation of inhibition that would reduce event specificity. Given that SST cells control size of memory engrams in the hippocampus (Stefanelli et al., 2016) as well as lateral inhibition in the auditory (Kato et al., 2017) and visual cortex (Adesnik et al., 2012), we reasoned that SST cells could control recruitment of cells during spontaneous activity. We selectively suppressed SST cells using a cre-dependent inhibitory hM4Di-DREADD during awake two-photon calcium imaging of spontaneous activity (Fig 5A). We tested the specificity of our construct through immunohistochemistry for somatostatin: 83 ± 8% of cells expressing the inhibitory hM4Di-DREADD also expressed somatostatin (n = 4 animals, Fig 5B) and 79 ± 6% of SST cells expressed the hM4Di-DREADD. We also confirmed that at this age, activation of the hM4Di-DREADD construct with clozapine reduced excitability of DREADD-expressing cells *in vitro* (Supplementary Fig 2A) similar to the excitability reducing effect of hM4Di-DREADD activation in developing layer 2/3 pyramidal neurons described previously (Naskar et al., 2019).

**Figure 4.**
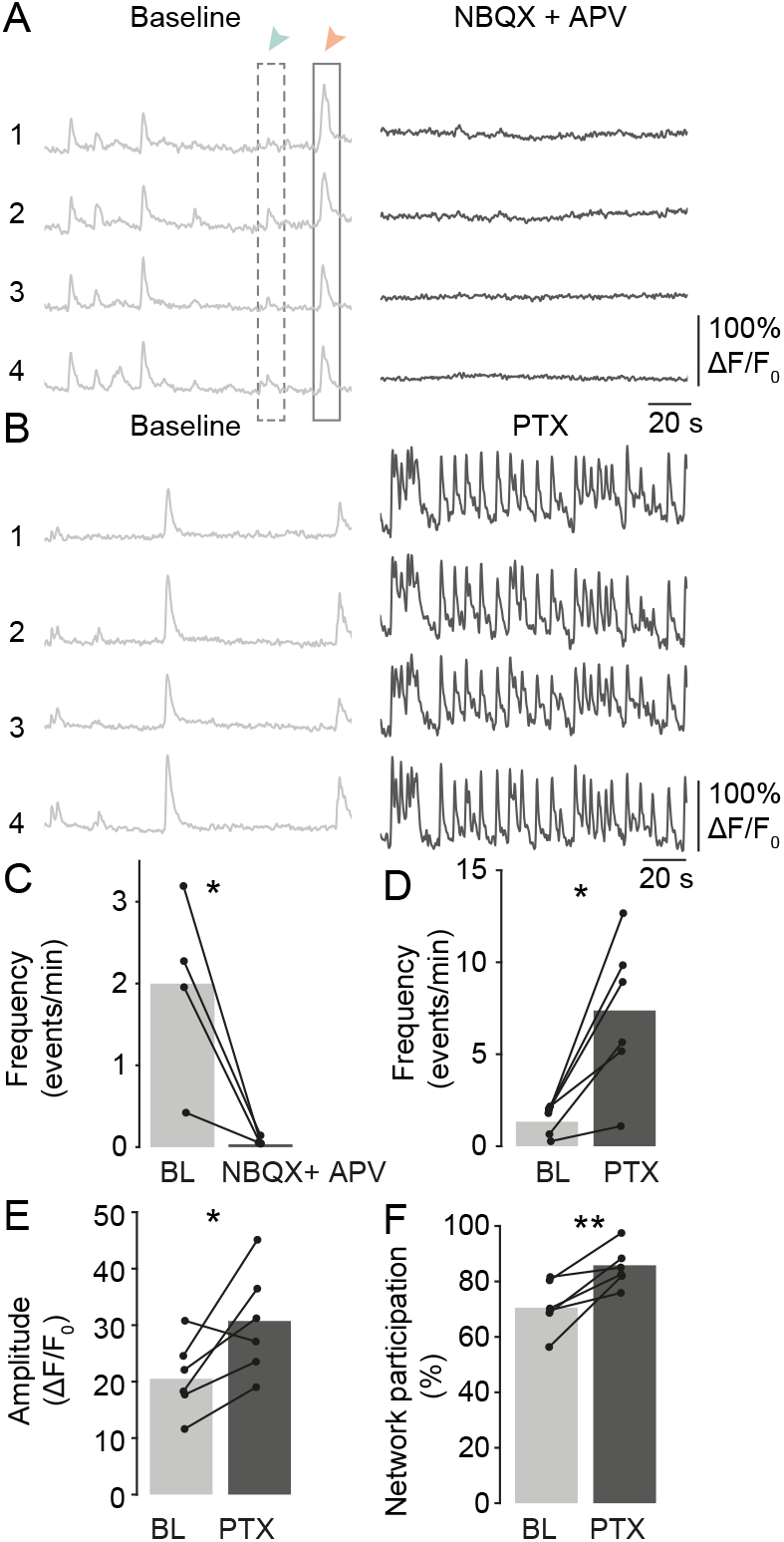
Both excitatory and inhibitory signaling are required for normal patterned spontaneous activity properties in the developing visual cortex A. Calcium traces from 4 cells before (left) and after (right) application of the AMPA receptor antagonist NBQX (50 μM) and the NMDA receptor antagonist APV (100 μM). B. Calcium traces from 4 cells before (left) and after (right) application of the GABAA receptor antagonist picrotoxin (330 μM). C. Blocking ionotropic glutamate receptors prevented almost all spontaneous network activity (p = 0.044, paired t-test, n = 4 animals). D. Blocking GABAergic signaling with PTX significantly increased the frequency of spontaneous activity (p = 0.01, paired t-test, n = 6 animals). E. Applying PTX significantly increased the network event amplitude (p = 0.04, paired t-test, n = 6 animals). F. The percentage of cells active in each event increased upon PTX application (p = 0.008, paired t-test, n = 6 animals).

For *in vivo* recordings, the fast-acting DREADD agonist clozapine (0.5 mg/kg) was subcutaneously injected to suppress SST cell activity. An example recording before and after the injection, including an average trace and participating cells, is shown in Fig 5C.

We found that upon SST cell suppression, the pairwise neuronal calcium trace correlation increased in animals expressing the hM4Di-DREADD construct (Supplementary Fig 2B, C), but not in control animals (Supplementary Fig 2D). In addition, suppressing SST cells specifically increased the frequency of events in the highest participation bin (Fig 5D). In control animals, there was no significant increase in frequency (Supplementary Fig 2E,F). Consequently, the increase observed in the highest participation bin of animals expressing the hM4Di-DREADD construct was significantly different from changes in control animals.

Two possible scenarios could underlie this increase in high participation events: either SST suppression allows more H-events to occur in the network or, alternatively, SST suppression causes L-events to recruit more cells, increasing participation. We did not detect an increase in mean event amplitude (Fig 5E, Supplementary Fig 2G), suggesting that neuronal action potential firing during events was unchanged. Furthermore, careful analysis of neuronal participation and amplitude revealed that clozapine specifically facilitated high participation events of low amplitude (Fig 5F, for controls lacking hM4Di-DREADD, see Supplementary Figure 2G). As H-events are characterized by their high amplitudes, these data suggest that SST neuron suppression selectively increased the number of cells participating in an L-event, rather than causing additional H-events. SST neuron suppression can therefore perturb some aspects of activity patterning, such as cell recruitment and pairwise correlation, while leaving others unaffected, like the firing rate of active neurons.

**Figure 5.**
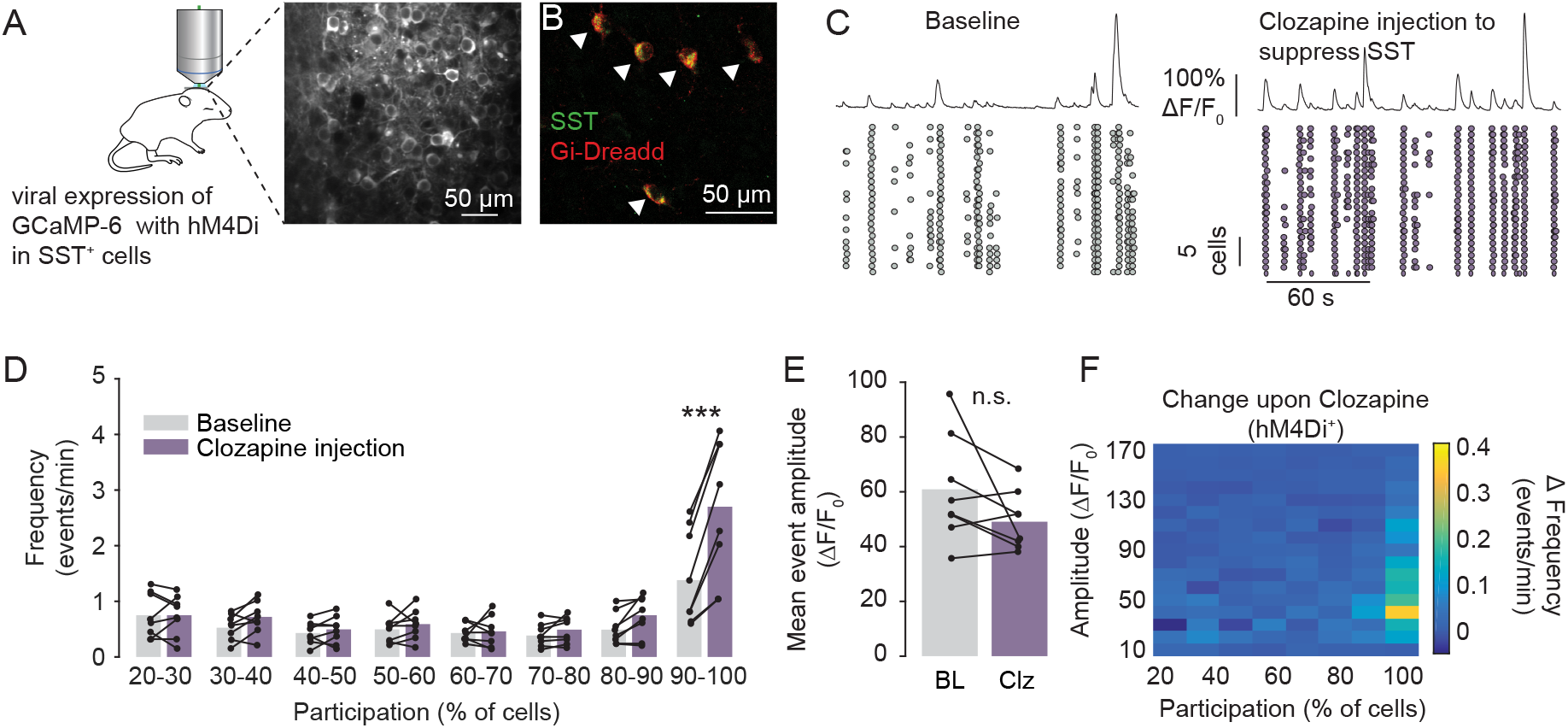
Suppressing somatostatin cell activity selectively increases the number of high-participation low-amplitude events A. Two-photon imaging was used to record calcium transients in animals expressing GCaMP-6s and an hM4Di construct in SST cells. Clozapine was injected at low concentrations to activate the hM4Di-DREADD and reduce cell excitability. B. Section of the visual cortex of a P9 SST-cre mouse injected with AAV-hSyn-DIO-hM4D(Gi)-mCherry (red) with immunostaining for somatostatin (green). All cells in the field of view were double-labeled. C. Neurons labelled with GCamP-6s via viral injection were imaged in awake pups during the second postnatal week. Example recording before (left) and after (right) injection of clozapine (DREADD agonist). Shown is the average network activity (above) and the activity of each imaged cell (below). Each circle represents a calcium transient in a cell. D. After clozapine injection in SST-cre mice expressing hM4Di, there is a significant increase in frequency of events in the highest participation bracket (p = 0.0002, threshold for significance at 0.0063 after Bonferroni correction for 8 comparisons, paired t-test, n = 8 animals). E. No change in amplitude was detected upon SST suppression using the DREADD agonist clozapine (n.s., paired t-test, n = 8 animals). BL = baseline, Clz = clozapine. F. Heat map showing the change in frequency of activity in events/min after injecting the DREADD agonist clozapine. The largest increase occurred in high-participation/low-am-plitude events.

### Somatostatin interneuron suppression increases spatial extent of network events

Next, we asked next how this control of cell recruitment affected the spatial extent of network activity. We used the same inhibitory hM4Di-DREADD construct to suppress SST cell activity whilst recording network activity across a large part of the cortex with wide field imaging (4.9 mm^2^). We first quantified activity origin and spread of all events (Fig 6A). Both before and after SST suppression, most activity originating in V1 was confined to this area (Fig 6B, C), indicating that spontaneous activity was restrained by V1 boundaries independently of SST neuron signaling. However, network events activated a significantly larger area of the cortex upon SST suppression (Fig 6D, E). This was due to both an increase in the lateral spread (Fig 6F) as well as the distance travelled during each event (Fig 6G). The duration of events did not change (Fig 6H), but their speed increased (Fig 6I) and allowed them to travel further. The event amplitude also increased slightly (Fig 6J), reflecting the increased participation of cells during each event given that there was no change in the amplitude of individual cells in two-photon imaging (Fig. 5E). The frequency of activity did not increase in the wide field imaging (Fig 6K), and instead showed a trend to decrease. The increase in frequency seen in 2-photon imaging therefore presumably reflected a larger proportion of events entering the field of view due to their lateral spread, rather than a true increase in the number of events. We found no significant changes upon clozapine administration to control animals (Supplementary Figure 3).

A histogram of event size (here measured as the area activated in a maximum projection of each event) before and after SST suppression showed a shift away from small events and towards larger events (Fig 6L). Given that the largest events did not show a change, we hypothesized that H-events were less affected by SST suppression than L-events, in line with the data shown in Fig 5. Using the same clustering method as in Fig 1, we split events into Land H-events and found a significant increase in mean activation area in L-events upon SST suppression (Fig 6M). H-events did not show an increase in area, possibly because excitation overrides SST lateral inhibition during H-events (Fig 6N). Together, the reduction of SST neuron activity restricts the lateral spread and cell activation density of L-events, but not their frequency or the overall firing rates of individual neurons during network events.

**Figure 6.**
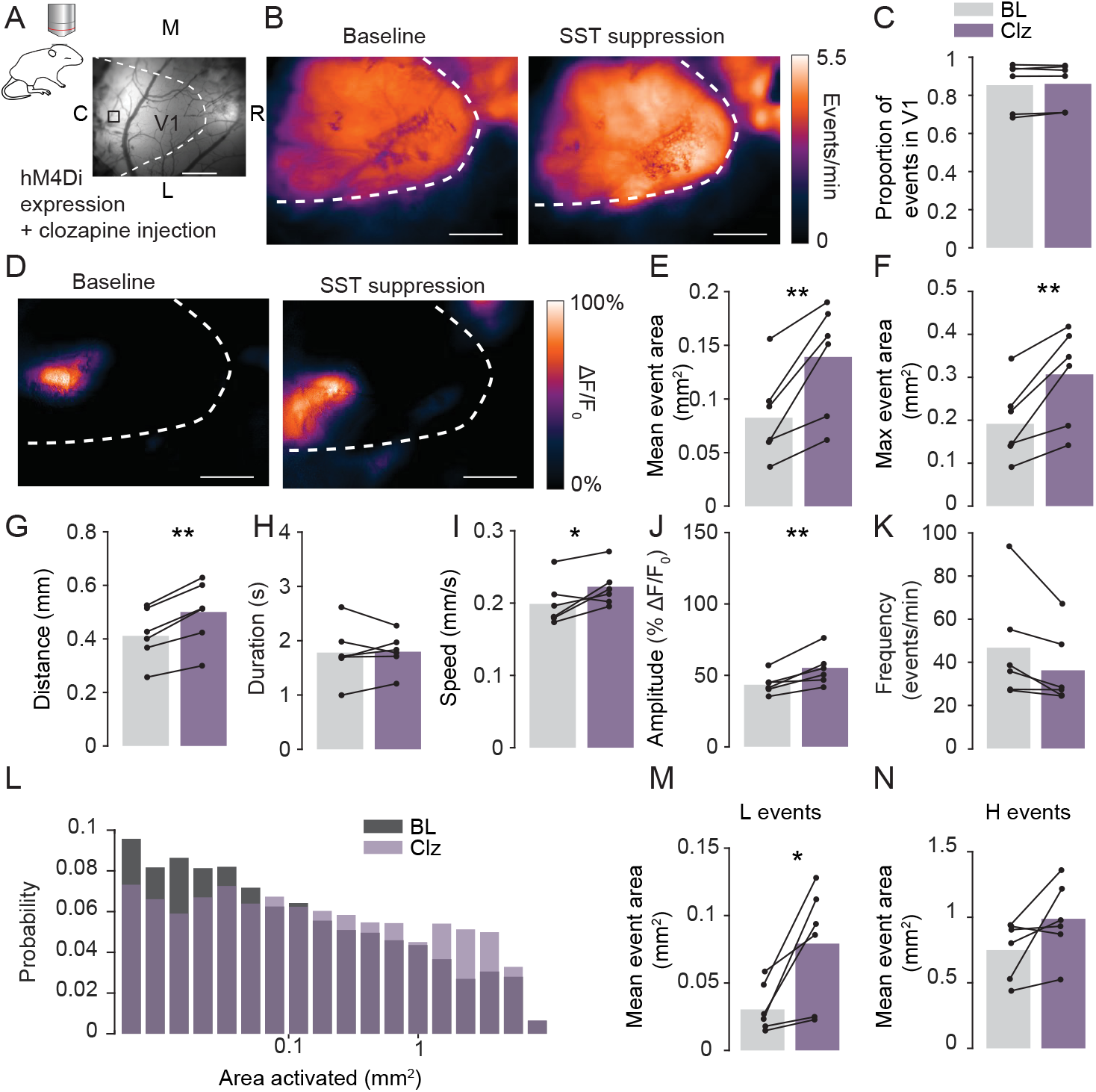
Somatostatin cell suppression selectively leads to larger event spread. A. Wide-field imaging was used to record large scale activity patterns in animals expressing GCaMP-6s and an hM4Di construct in SST cells (M: medial, L: lateral, R: rostral, C: caudal). The hM4Di-DREADD was activated with clozapine (Clz) injection. B. Average activity frequency per pixel before (left) and after (right) clozapine injection to induce SST suppression. C. There was no significant change in the proportion of events that activated V1 upon clozapine injection to induce SST suppression (n.s., paired t-test, n = 6 animals). D. Example event before (left) and after (right) SST suppression. E. The mean area activated by each event increased upon suppression of SST cells (p = 0.006, paired t-test, n = 6 animals). F. The max event area (the activated area during the largest frame of an event) was significantly higher after SST suppression (p = 0.006, paired t-test, n = 6 animals). G. Events traveled further across the cortex after suppression of SST cells through clozapine injection (p = 0.00066, paired t-test, n = 6 animals). H. After SST suppression, the duration of events did not change (n.s, paired t-test, n = 6 animals). I. The speed of events increased after SST suppresion (p = 0.049, paired t-test, n = 6 animals). J. The mean event amplitude increased after SST suppression (p = 0.0095, paired t-test, n = 6 animals). K. The frequency of events did not change upon suppression of SST (n.s., paired t-test, n = 6 animals). L. Histogram of the area activated by a maximum projection of all frames of each event, showing a shift from smaller to larger areas. M. L-event area increased significantly after SST suppression (p = 0.03, paired t-test, n = 6 animals). N. H-event area did not change significantly after SST suppression (n.s., paired t-test, n = 6 animals).

## Discussion

We show here that inhibitory control is required for normal activity features as early as the second postnatal week. Furthermore, the balance between excitatory and inhibitory signaling determines those characteristics of spontaneous activity patterns that convey information about the structure and connectivity of the developing visual system.

A large body of work has shown that the features of spontaneous activity patterns are instructive in wiring the developing brain (Feller, 1999; Katz and Shatz, 1996). A particularly well investigated model is the visual system, where manipulations of the size or frequency of retinal waves have directly demonstrated the importance of accurate patterning of spontaneous activity for fine-tuning the ascending visual pathways (Burbridge et al., 2014; Xu et al., 2015). If activity is too highly correlated, a broader set of input cells will be coincidently active with the postsynaptic cell, preventing Hebbian-based mechanisms from distinguishing between synaptic connections that should be stabilized and those that should be eliminated (Kirkby et al., 2013). Using a wide range of techniques, we have described L-events as activity patterns during which relatively few cells are weakly activated. We hypothesize that the largely retinal origin (Siegel et al., 2012) and restricted size of L-events allows them to mediate refinement of the network, as their localized nature could maintain topographic specificity. In contrast, H-events are highly synchronized in time and drive substantial areas of the cortex to fire large numbers of action potentials. The high spatial spread and temporal synchronicity of H-events makes them less suitable to convey information about cell arrangement in the retina. Instead, H-events may perform synaptic homeostasis to maintain workable ranges of synaptic strength, perhaps in a similar mechanism as during slow-wave sleep (González-Rueda et al., 2018; Tononi and Cirelli, 2006). In the present study, we found that the E/I ratio is increased during H-events compared to L-events. Accordingly and consistent with previous studies, blocking inhibition pharmacologically increased activity levels (Kirmse et al., 2015; Minlebaev et al., 2007) and pushed spontaneous activity patterns towards H-like characteristics, essentially eliminating exactly those features of L-type patterns that most likely drive synaptic refinement. Therefore, we conclude that inhibition is required for shaping activity patterns that mediate precise connectivity in the developing visual cortex.

We observed significant increases of the E/I ratio during H-compared to L-events on the level of synaptic inputs as well as the output of excitatory versus inhibitory neurons. Quantitatively, these increases differed, however, and were quite high for synaptic inputs, but relatively small for the outputs. This difference could be a consequence of the immature state of the interneurons at this age. Given that voltage-clamp experiments are spatially limited, it seems likely that we are primarily recording inputs from soma-targeting parvalbumin-positive basket cells rather than, for instance, dendrite-targeting SST cells. Basket cells do not express the calcium binding protein PV until a sudden onset after P13 (Gonchar et al., 2008). As the excitatory/inhibitory ratio was higher during H-events, we hypothesize that these basket cells can restrict firing frequency during both L- and H-events, but do not increase their output sufficiently during H-events to keep up with excitation. The calcium-buffering properties of parvalbumin are not required for the ability of PV neurons to fire at high frequencies, but are important in maintaining synaptic strength during fast-spiking, non-adaptive firing (Schwaller et al., 2002). Therefore, we speculate that once PV is expressed at P14, PV synapses may be able to keep up even during strong excitation. Indeed, the strength of the synaptic connection between fast spiking interneurons and pyramidal cells in the visual cortex was recently found to increase significantly during eye opening (Guan et al., 2017). More powerful transmission at inhibitory synapses could reduce or prevent the occurrence of H-events, contributing to the progressive desynchronization of spontaneous activity around eye-opening (Rochefort et al., 2009). In addition, this maturation step may underlie the increase of visually driven inhibitory responses that facilitate the emergence of up-states during wakefulness (Colonnese, 2014) and eventually determine the onset of the critical period for ocular dominance (Fagiolini and Hensch, 2000) when preferential suppression of spontaneous activity may increase the relative contribution of visually evoked activity to refinement (Toyoizumi et al., 2013).

Intriguingly, specifically the density of activation and the lateral spread of events, rather than other event characteristics, could be manipulated by suppressing SST cells. This could be due to the reduction of the excitatory/inhibitory balance, or due to specific actions of SST cells. A specific action would match well with the role of SST in the adult, where SST suppression increases cell recruitment in hippocampal engrams (Stefanelli et al., 2016) and increases network synchronicity (Chen et al., 2015). SST cells have extensive axonal arbors in L1 (Urban-Ciecko and Barth, 2016) and excellent control over large distances through lateral inhibition in the visual (Adesnik et al., 2012) and auditory (Kato et al., 2017) cortex. Even in the young cortex, SST neurons perform - transient - network functions (Marques-Smith et al., 2016; Tuncdemir et al., 2016). Since the spread of activity patterns constrains the degree of synaptic refinement, SST neurons may optimize early activity patterns for establishing precise neuronal connections during the second postnatal week. It is possible that SST neurons may have an excitatory role in the even younger cortex (P1-5) as reported recently for the neonatal hippocampus (Flossmann et al., 2019).

In conclusion, we find that interneurons exert robust inhibitory control over network activity during the second postnatal week, even though inhibitory signaling is not fully strengthened until after eye opening. Furthermore, the different balance between excitation and inhibition during L- and H-events indicates that acute modulation of inhibition is essential in shaping the features of spontaneous activity. It seems that not only do interneurons exert inhibitory control over this early age, but they delicately shape activity patterns crucial for driving fine-tuning of the developing network.

## Methods

### Animals

All experimental procedures were approved by the institutional animal care and use committee of the Royal Netherlands Academy of Sciences. Mice of both sexes were used. All animals were aged between postnatal days (P) 8-14. Wildtype mice were either C57BL/6J mice or C57BL/6J x CBA F1. These mice open their eyes at P14. Gad2 x tdTomato mice were generated by crossing the reporter CAG-tdTomato mice line (td, Jackson labs) with GAD-Cre (GAD2-IRES-Cre, Jackson labs). SST-cre (JAX 13044) and VIP-cre (JAX 10908) mice were designed by Dr. Z. Josh Huang and ordered from Jackson Labs (SST-IRES-Cre; Taniguchi et al., 2011.)

### Surgery

Animals were anesthetized with isoflurane (3% in 1 l/min O_2_). After anesthesia had become effective, lidocaine was used for local analgesia and a head bar with an opening (Ø 4 mm) above the visual cortex (0.5-2.5 mm rostral from lambda and 1-3 mm lateral from the midline) was attached to the skull with superglue and dental cement. For calcium imaging, a small craniotomy above the visual cortex was performed. The exposed cortical surface was kept moist with cortex buffer (125 mM NaCl, 5 mM KCl, 10 mM glucose, 10 mM HEPES, 2 mM MgSO4 and 2 mM CaCl2 [pH 7.4]). For calcium imaging under anesthesia, isoflurane levels were lowered to 0.7-1%. Prior to awake imaging, the animals were given 60 minutes to recover from anesthesia.

### Virus production

AAV vectors were produced as described previously (Verhaagen et al., 2018). In short, AAV1 serotype helper plasmid and pAAV-hSyn-DIO-hM4D(Gi)-mCherry or pAAV-hSyn-GCaMP6s were co-transfected into HEK293T cells. Seventy-two hours later, the cells were harvested, lysed and centrifuged. Subsequently the viral particles were purified from the supernatant using an iodixanol density gradient and further concentrated using an Amicon Ultra-15 centrifugal filter. Titers were determined by quantitative PCR for the WPRE element in the viral genomes (vg). Primers for WPRE: 5’-CCCACTTGGCAGTACATCAA-3’ and 5’-GGAAAGTCCCATAAGGTCATGT-3’. Titers were 2E+12 vg/ml for pAAV-hSyn-DIO-hM4D(Gi)-mCherry and pAAV-hSyn-GCAMP6s.

### Virus injection

Pups (P0-P1) were anaesthetized using hypothermia (6 minutes). A small cut was made in the skin before insertion of a glass pipette and injection of 27 nl of virus in V1 (two-photon imaging) or V1 and RL (wide field imaging). Animals were injected with a mix of 1:1 AAV1-hSyn-DIO-hM4D(Gi)-mCherry and AAV1-hSyn-GCaMP6s or a mixture of 1:1 PBS and AAV1-hSyn-GCaMP6s. pAAV-hSyn-DIO-hM4D(Gi)-mCherry was a gift from Bryan Roth (Krashes et al., 2011, Addgene plasmid # 44362).

### In Utero Electroporation

For wide-field calcium imaging shown in Figure 1, pyramidal neurons in layer 2/3 of the visual cortex were transfected with GCaMP6s (2 mg/ml) and DsRed (2 mg/ml) at E16.5 using in utero electroporation (Harvey et al., 2009). Pregnant mice were anesthetized with isoflurane and a small incision (1.5-2 cm) was made in the abdominal wall. The uterine horns were carefully removed from the abdomen, and DNA was injected into the lateral ventricle of embryos using a sharp glass electrode. Voltage pulses (five square wave pulses, 30 V, 50-ms duration, 950-ms interval, custom-built electroporator) were delivered across the brain with tweezer electrodes covered in conductive gel. Embryos were rinsed with warm saline solution and returned to the abdomen, after which time the muscle and skin were sutured.

### Calcium indicator application

The calcium-sensitive dye Oregon Green 488 BAPTA-1 AM (OGB-1, Invitrogen) was dissolved in 4 μl pluronic F-127, 20% solution in DMSO (Invitrogen) and further diluted (1: 10) in dye buffer (150 mM NaCl, 2.5 mM KCl, and 10 mM HEPES) to yield a final concentration of 1 mM. The dye was then pressure-ejected at 10-12 psi for 12-13 min with a micropipette (3-5 MΩ) attached to a picospritzer (Toohey). As OGB-1AM has a linear relationship between fluorescence and the number of action potentials fired in both pyramidal cells and PV cells, the most common GABAergic interneuron (Hofer et al., 2011; Kerlin et al., 2010), we used calcium amplitude as an approximation of cell firing.

### DREADD activation

Animals expressing the above virus were first imaged at baseline. The fast DREADD agonist clozapine was then injected subcutaneously at 0.5 mg/kg. We expected to see effects in a relatively narrow time window and therefore restricted our analyses to 30 minutes postinjection.

### Image acquisition

*In vivo* calcium imaging was performed on either a Nikon (A1R-MP) with a 0.8/16x water-immersion objective and a Ti:Sapphire laser (Chameleon II, Coherent) or a Movable Objective Microscope (Sutter Instruments) with a Ti:Sapphire laser (MaiTai HP, Spectra Physics) and a 0.8/40x water-immersion objective (Olympus) using Nikon or ScanImage software (Pologruto et al., 2003). We recorded the movement signal of the scan mirrors to synchronize calcium imaging and electrophysiology. Pixel size was 300 nm and images of 330 by 330 μm were recorded at 5-10 Hz.

Epifluorescence was recorded using custom-built LabVIEW software (National Instruments) using a digital CCD camera (QImaging), a 0.16/4x (Olympus) objective and a xenon-arc lamp (Sutter Instrument Company).

### In vivo whole-cell and extracellular electrophysiology

Membrane potential was recorded in current clamp at 10 kHz and filtered at 3 kHz (Multiclamp 700b; Molecular Devices). For current clamp recordings electrodes (4.5-6 MΩ) were filled with intracellular solution (105 mM K gluconate, 10 mM HEPES, 30 mM KCl, 10 mM phosphocreatine, 4 mM MgATP, and 0.3 mM GTP; Golshani et al., 2009). 10 μM Alexa 594 hydrazide (Invitrogen) was added to allow targeted whole-cell recordings. The mean network participation of these events based on calcium imaging was 45% (L-events) and 94% (H-events). The patched cells participated (fired at least 1 action potential) in 70% of L-events and 100% of H-events. Taking a threshold of three action potentials or more gives participation rates similar to the calcium imaging (41% and 94%), implying that we can pick up cells that fire three action potentials or more. No correction was made for the liquid junction potential.

Synaptic currents were recorded in voltage clamp at 10 kHz and filtered at 3 kHz (Multiclamp 700b; Molecular Devices). For voltage clamp recordings, electrodes were filled with intracellular solution (120 mM CsMeSO_3_, 8 mM NaCl, 15 mM CsCl_2_, 10 mM TEA-Cl, 10 mM HEPES, 5 mM QX-314, 4 mM MgATP, 0.3 mM Na-GTP, Kwon et al., 2012).

Raw unfiltered local field potentials (LFP) were measured with a glass pipette with resistance (<4 MΩ). The serial resistance was monitored during the recording and recordings made with a resistance higher than 12 MΩ were excluded from the data.

### Image processing

Two-photon image processing: To remove drift and movement artifacts from each recording, we performed image alignment using NoRMCorre (Pnevmatikakis and Giovannucci, 2017). Each recording was aligned to the first recording in the series to remove any movements between recording sessions. Delta F stacks were made using the average fluorescence per pixel as baseline. ROIs were hand-drawn using ImageJ (NIH). Automated transient detection and further data processing was performed using custom-made Matlab software (MathWorks).

Epifluorescence image processing: Delta F stacks were made using the average fluorescence per pixel as baseline. V1 was identified based on activity coordinates and shape after this method of identification was confirmed through immunohistochemistry for vGluT2.

### Data analysis and statistics

Electrophysiology measurements were aligned to images using custom-built Matlab software. The LFP was bandpass-filtered offline, (3-100 Hz, 10th order Buttersworth filter). The frequency components were analyzed with the open source MATLAB library Chronux (Bokil et al., 2010). Hierarchical clustering, silhouette analysis, receiver-operation characteristic curves and random forest classification were analyzed in Matlab. One animal was excluded from random forest classification due to insufficient inhibitory events to allow for good out-of-bag measurements. As described in Montijn et al., (2016), we used Ward’s method (Ward, 1963) hierarchical clustering to construct dendrograms of all events. Clustering was based on the number of action potentials fired and duration (Fig 1F) or the amplitude and activated area (when performed on wide-field data). Hierarchical clustering was performed within each animal. Silhouette curves were made based on the dendrogram (Rousseeuw, 1987), where a single datapoint in a cluster was given a silhouette value of 0. The optimal number of clusters was taken as the overall maximum.

In all but one statistical test, we used paired measurements from each animal and therefore performed paired two-tailed paired t-tests. When DREADD-expressing and non-DREADD expressing animals were compared, a two-sample two-tailed t-test was used. A Bonferroni correction for multiple comparisons was used where more than one test was performed.

When the distribution of inter-spike intervals was compared between L- and H-events in Fig 1D, a two-sample Kolmogorov-Smirnov test was used. Pairwise correlation between cell calcium activity (Figure 5) was calculated using Spearman’s rank-order correlation on the entire trace after delta F calculation.

### *In vitro* whole-cell patch-clamp recordings

Acute 300 μm coronal slices of the visual cortex were dissected. Pups were sacrificed by decapitation and their brains were immersed in ice-cold cutting solution (in mM): 2.5 KCl, 1.25 NaH_2_PO_4_, 26 NaHCO_3_, 20 Glucose, 215 Sucrose, 1 CaCl_2_, 7 MgCl_2_ (Sigma), pH 7.3-7.4, bubbled with 95%/5% O_2_/CO_2_. Slices were obtained with a vibratome (VT1200 S, Leica) and subsequently incubated at 34°C in artificial cerebrospinal fluid (ACSF, in mM): 125 NaCl, 3.5 KCl, 1.25 NaH_2_PO_4_, 26 NaHCO_3_, 20 Glucose, 2 CaCl_2_, 1 MgCl2 (Sigma), pH 7.3-7.4. After 45 minutes, slices were transferred to the electrophysiology setup, kept at room temperature and bubbled with 95%/5% O_2_/CO_2_. For patch recordings, slices were transferred to a recording chamber and perfused (3 ml/min) with ACSF solution bubbled with 95%/5% O_2_/CO_2_ at 34°C.

Layer 2/3 SST^+^ or SST^-^ cells were identified using an fluorescence/IR-DIC video microscope (Olympus BX51WI). SST^+^ interneurons were identified by the mCherrry protein fluorescence from mice injected with pAAV-hSyn-DIO-hM4D(Gi)-mCherry. Current-clamp recordings were made with a MultiClamp 700B amplifier (Molecular Devices), filtered with a low pass Bessel filter at 10 kHz and digitized at 20-50 kHz (Digidata 1440A, Molecular Devices). Series resistance was assessed during recordings and neurons showing a series resistance > 30 MΩ or change > 30% were discarded. Digitized data were analyzed offline using Clampfit 10 (Molecular Devices) and Igor (WaveMetrics).

Electrodes were filled with an intracellular solution containing (in mM): 122 KGluconate, 10 Hepes, 13 KCl, 10 phosphocreatine disodium salt hydrate, 4 ATP magnesium salt, 0.3 GTP sodium salt hydrate (Sigma), pH 7.3. Clozapine 10 μM (Tocris) was bath applied.

After breaking the seal, variable current injection was applied to keep cells at −60 mV. To test the excitability of the cells, current injection from −80 pA to 160 pA was applied in 10 pA increments.

### Analysis of in vitro electrophysiological experiments

Input-output curves were generated by calculating the cell firing rate at each current injection step in control and clozapine conditions.

## Acknowledgements

The authors thank Cátia Silva, Christiaan Levelt, Alexander Heimel and Julijana Gjorgjieva for their comments on this manuscript, Monique van Mourik for technical assistance and Johan Winnubst for help with data analysis software. This work was supported by grants of the Netherlands Organization for Scientific Research (NWO, ALW Open Program grants, no. 822.02.006 and ALWOP.216; ALW Vici grant, no. 865.12.001) and the “Stichting Vrienden van het Herseninstituut”.

## Declaration of Interests

The authors declare no competing interests.

## References

Ackman, J.B., and Crair, M.C. (2014). Role of emergent neural activity in visual map development. Curr. Opin. Neurobiol. 24, 166–175.

Ackman, J.B., Burbridge, T.J., and Crair, M.C. (2012). Retinal waves coordinate patterned activity throughout the developing visual system. Nature 490, 219–225.

Adesnik, H., Bruns, W., Taniguchi, H., Huang, Z.J., and Scanziani, M. (2012). A neural circuit for spatial summation in visual cortex. Nature 490, 226–231.

Allene, C., and Cossart, R. (2010). Early NMDA receptor-driven waves of activity in the developing neocortex: physiological or pathological network oscillations? J.Physiol 588, 83–91.

Blankenship, A.G., and Feller, M.B. (2010). Mechanisms underlying spontaneous patterned activity in developing neural circuits. Nat. Rev. Neurosci. 11, 18–29.

Bokil, H., Andrews, P., Kulkarni, J.E., Mehta, S., and Mitra, P. (2010). Chronux: A Platform for Analyzing Neural Signals. J. Neurosci. Methods 192, 146–151.

Breiman, L. (2001). Random Forests. Mach. Learn. 45, 5–32.

Burbridge, T.J., Xu, H.-P., Ackman, J.B., Ge, X., Zhang, Y., Ye, M.-J., Zhou, Z.J., Xu, J., Contractor, A., and Crair, M.C. (2014). Visual Circuit Development Requires Patterned Activity Mediated by Retinal Acetylcholine Receptors. Neuron 84, 1049–1064.

Cang, J., Renteria, R.C., Kaneko, M., Liu, X., Copenhagen, D.R., and Stryker, M.P. (2005). Development of Precise Maps in Visual Cortex Requires Patterned Spontaneous Activity in the Retina. Neuron 48, 797–809.

Che, A., Babij, R., Iannone, A.F., Fetcho, R.N., Ferrer, M., Liston, C., Fishell, G., and García, N.V.D.M. (2018). Layer I Interneurons Sharpen Sensory Maps during Neonatal Development. Neuron 99, 98–116.

Chen, N., Sugihara, H., and Sur, M. (2015). An acetylcholine-activated microcircuit drives temporal dynamics of cortical activity. Nat. Neurosci. 18, 892–902.

Cherubini, E., Gaiarsa, J.L., and Ben-Ari, Y. (1991). GABA: an excitatory transmitter in early postnatal life. Trends Neurosci. 14, 515–519.

Cline, H. (2003). Sperry and Hebb: oil and vinegar? Trends Neurosci. 26, 655–661.

Colonnese, M.T. (2014). Rapid Developmental Emergence of Stable Depolarization during Wakefulness by Inhibitory Balancing of Cortical Network Excitability. J. Neurosci. 34, 5477–5485.

Colonnese, M.T., and Phillips, M.A. (2018). Thalamocortical function in developing sensory circuits. Curr. Opin. Neurobiol. 52, 72–79.

Duan, Z.R.S., Che, A., Chu, P., Modol, L., Bollmann, Y., Babij, R., Fetcho, R.N., Otsuka, T., Fuccillo, M.V., Liston, C., et al. (2019). GABAergic Restriction of Network Dynamics Regulates Interneuron Survival in the Developing Cortex. Neuron 0.

Fagiolini, M., and Hensch, T.K. (2000). Inhibitory threshold for critical-period activation in primary visual cortex. Nature 404, 183–186.

Feller, M.B. (1999). Spontaneous correlated activity in developing neural circuits. Neuron 22, 653–656.

Flossmann, T., Kaas, T., Rahmati, V., Kiebel, S.J., Witte, O.W., Holthoff, K., and Kirmse, K. (2019). Somatostatin Interneurons Promote Neuronal Synchrony in the Neonatal Hippocampus. Cell Rep. 26, 3173–3182.e5.

Golshani, P., Goncalves, J.T., Khoshkhoo, S., Mostany, R., Smirnakis, S., and Portera-Cailliau, C. (2009). Internally Mediated Developmental Desynchronization of Neocortical Network Activity. J. Neurosci. 29, 10890–10899.

Gonchar, Y., Wang, Q., and Burkhalter, A.H. (2008). Multiple distinct subtypes of GABAergic neurons in mouse visual cortex identified by triple immunostaining. Front. Neuroanat. 1, 3.

González-Rueda, A., Pedrosa, V., Feord, R.C., Clopath, C., and Paulsen, O. (2018). Activity Dependent Downscaling of Subthreshold Synaptic Inputs during Slow-Wave-Sleep-like Activity In Vivo. Neuron 97, 1244–1252.

Gribizis, A., Ge, X., Daigle, T.L., Ackman, J.B., Zeng, H., Lee, D., and Crair, M.C. (2019). Visual Cortex Gains Independence from Peripheral Drive before Eye Opening. Neuron 0.

Guan, W., Cao, J.-W., Liu, L.-Y., Zhao, Z.-H., Fu, Y., and Yu, Y.-C. (2017). Eye opening differentially modulates inhibitory synaptic transmission in the developing visual cortex. ELife 6, e32337.

Hanganu, I.L., Ben Ari, Y., and Khazipov, R. (2006). Retinal waves trigger spindle bursts in the neonatal rat visual cortex. J Neurosci 26, 6728–6736.

Harris, K.D., and Thiele, A. (2011). Cortical state and attention. Nat. Rev. Neurosci. 12, 509.

Hofer, S.B., Ko, H., Pichler, B., Vogelstein, J., Ros, H., Zeng, H., Lein, E., Lesica, N.A., and Mrsic-Flogel, T.D. (2011). Differential connectivity and response dynamics of excitatory and inhibitory neurons in visual cortex. Nat Neurosci 14, 1045–1052.

Kato, H.K., Asinof, S.K., and Isaacson, J.S. (2017). Network-Level Control of Frequency Tuning in Auditory Cortex. Neuron 95, 412–423.

Katz, L.C., and Shatz, C.J. (1996). Synaptic activity and the construction of cortical circuits. Science 274, 1133–1138.

Kepecs, A., and Fishell, G. (2014). Interneuron Cell Types: Fit to form and formed to fit. Nature 505, 318–326.

Kerlin, A.M., Andermann, M.L., Berezovskii, V.K., and Reid, R.C. (2010). Broadly tuned response properties of diverse inhibitory neuron subtypes in mouse visual cortex. Neuron 67, 858–871.

Kerschensteiner, D. (2014). Spontaneous Network Activity and Synaptic Development. Neurosci. Rev. J. Bringing Neurobiol. Neurol. Psychiatry 20, 272–290.

Khazipov, R., Sirota, A., Leinekugel, X., Holmes, G.L., Ben-Ari, Y., and Buzsaki, G. (2004). Early motor activity drives spindle bursts in the developing somatosensory cortex. Nature 432, 758–761.

Kirkby, L.A., Sack, G.S., Firl, A., and Feller, M.B. (2013). A Role for Correlated Spontaneous Activity in the Assembly of Neural Circuits. Neuron 80, 1129–1144.

Kirmse, K., Kummer, M., Kovalchuk, Y., Witte, O.W., Garaschuk, O., and Holthoff, K. (2015). GABA depolarizes immature neurons and inhibits network activity in the neonatal neocortex in vivo. Nat. Commun. 6, 7750.

Ko, H., Cossell, L., Baragli, C., Antolik, J., Clopath, C., Hofer, S.B., and Mrsic-Flogel, T.D. (2013). The emergence of functional microcircuits in visual cortex. Nature 496, 96–100.

Krashes, M.J., Koda, S., Ye, C., Rogan, S.C., Adams, A.C., Cusher, D.S., Maratos-Flier, E., Roth, B.L., and Lowell, B.B. (2011). Rapid, reversible activation of AgRP neurons drives feeding behavior in mice. J. Clin. Invest. 121, 1424–1428.

Leighton, A.H., and Lohmann, C. (2016). The Wiring of Developing Sensory Circuits-From Patterned Spontaneous Activity to Synaptic Plasticity Mechanisms. Front. Neural Circuits 10, 71.

Luhmann, H.J., and Khazipov, R. (2018). Neuronal Activity Patterns in the Developing Barrel Cortex. Neuroscience 368, 256–267.

Markram, H., Toledo-Rodriguez, M., Wang, Y., Gupta, A., Silberberg, G., and Wu, C. (2004). Interneurons of the neocortical inhibitory system. Nat.Rev.Neurosci 5, 793–807.

Marques-Smith, A., Lyngholm, D., Kaufmann, A.-K., Stacey, J.A., Hoerder-Suabedissen, A., Becker, E.B.E., Wilson, M.C., Molnár, Z., and Butt, S.J.B. (2016). A Transient Translaminar GABAergic Interneuron Circuit Connects Thalamocortical Recipient Layers in Neonatal Somatosensory Cortex. Neuron 89, 536–549.

Minlebaev, M., Ben-Ari, Y., and Khazipov, R. (2007). Network mechanisms of spindle-burst oscillations in the neonatal rat barrel cortex in vivo. J. Neurophysiol. 97, 692–700.

Modol, L., Bollmann, Y., Tressard, T., Baude, A., Che, A., Duan, Z.R.S., Babij, R., García, N.V.D.M., and Cossart, R. (2019). Assemblies of Perisomatic GABAergic Neurons in the Developing Barrel Cortex. Neuron 0.

Montijn, J.S., Olcese, U., and Pennartz, C.M.A. (2016). Visual Stimulus Detection Correlates with the Consistency of Temporal Sequences within Stereotyped Events of V1 Neuronal Population Activity. J. Neurosci. 36, 8624–8640.

Naskar, S., Narducci, R., Balzani, E., Cwetsch, A.W., Tucci, V., and Cancedda, L. (2019). The development of synaptic transmission is time-locked to early social behaviors in rats. Nat. Commun. 10, 1195.

Oh, W.C., Lutzu, S., Castillo, P.E., and Kwon, H.-B. (2016). De novo synaptogenesis induced by GABA in the developing mouse cortex. Science 353, 1037–1040.

Pnevmatikakis, E.A., and Giovannucci, A. (2017). NoRMCorre: An online algorithm for piecewise rigid motion correction of calcium imaging data. J. Neurosci. Methods 291, 83–94.

Pologruto, T.A., Sabatini, B.L., and Svoboda, K. (2003). ScanImage: flexible software for operating laser scanning microscopes. BiomedEng Online 2, 13.

Rochefort, N.L., Garaschuk, O., Milos, R.I., Narushima, M., Marandi, N., Pichler, B., Kovalchuk, Y., and Konnerth, A. (2009). Sparsification of neuronal activity in the visual cortex at eye-opening. Proc.Natl.Acad.Sci.U.S.A 106, 15049–15054.

Rochefort, N.L., Narushima, M., Grienberger, C., Marandi, N., Hill, D.N., and Konnerth, A. (2011). Development of direction selectivity in mouse cortical neurons. Neuron 71, 425–432.

Rousseeuw, P.J. (1987). Silhouettes: A graphical aid to the interpretation and validation of cluster analysis. J. Comput. Appl. Math. 20, 53–65.

Sanes, J.R., and Yamagata, M. (2009). Many paths to synaptic specificity. Annu. Cell DevBiol 25, 161–195.

Schwaller, B., Meyer, M., and Schiffmann, S. (2002). “New” functions for “old” proteins: The role of the calcium-binding proteins calbindin D-28k, calretinin and parvalbumin, in cerebellar physiology. Studies with knockout mice. The Cerebellum 1, 241–258.

Siegel, F., Heimel, J.A., Peters, J., and Lohmann, C. (2012). Peripheral and central inputs shape network dynamics in the developing visual cortex in vivo. Curr. Biol. 22, 253–258.

Stefanelli, T., Bertollini, C., Lüscher, C., Muller, D., and Mendez, P. (2016). Hippocampal Somatostatin Interneurons Control the Size of Neuronal Memory Ensembles. Neuron 89, 1074–1085.

Taniguchi, H., He, M., Wu, P., Kim, S., Paik, R., Sugino, K., Kvitsani, D., Fu, Y., Lu, J., Lin, Y., et al. (2011). A resource of cre driver lines for genetic targeting of GABAergic neurons in cerebral cortex. Neuron 71, 995–1013.

Tononi, G., and Cirelli, C. (2006). Sleep function and synaptic homeostasis. Sleep MedRev 10, 49–62.

Torborg, C.L., and Feller, M.B. (2005). Spontaneous patterned retinal activity and the refinement of retinal projections. Prog.Neurobiol. 76, 213–235.

Toyoizumi, T., Miyamoto, H., Yazaki-Sugiyama, Y., Atapour, N., Hensch, T.K., and Miller, K.D. (2013). A Theory of the Transition to Critical Period Plasticity: Inhibition Selectively Suppresses Spontaneous Activity. Neuron 80, 51–63.

Tremblay, R., Lee, S., and Rudy, B. (2016). GABAergic Interneurons in the Neocortex: From Cellular Properties to Circuits. Neuron 91, 260–292.

Tuncdemir, S.N., Wamsley, B., Stam, F.J., Osakada, F., Goulding, M., Callaway, E.M., Rudy, B., and Fishell, G. (2016). Early Somatostatin Interneuron Connectivity Mediates the Maturation of Deep Layer Cortical Circuits. Neuron 89, 521–535.

Urban-Ciecko, J., and Barth, A.L. (2016). Somatostatin-expressing neurons in cortical networks. Nat. Rev. Neurosci. 17, 401–409.

Valeeva, G., Abdullin, A., Tyzio, R., Skorinkin, A., Nikolski, E., Ben-Ari, Y., and Khazipov, R. (2010). Temporal coding at the immature depolarizing GABAergic synapse. Front Cell Neurosci 4.

Valeeva, G., Tressard, T., Mukhtarov, M., Baude, A., and Khazipov, R. (2016). An Optogenetic Approach for Investigation of Excitatory and Inhibitory Network GABA Actions in Mice Expressing Channelrhodopsin-2 in GABAergic Neurons. J. Neurosci. 36, 5961–5973.

Verhaagen, J., Hobo, B., Ehlert, E.M.E., Eggers, R., Korecka, J.A., Hoyng, S.A., Attwell, C.L., Harvey, A.R., and Mason, M.R.J. (2018). Small Scale Production of Recombinant Adeno-Associated Viral Vectors for Gene Delivery to the Nervous System. Methods Mol. Biol. Clifton NJ 1715, 3–17.

van Versendaal, D., and Levelt, C.N. (2016). Inhibitory interneurons in visual cortical plasticity. Cell. Mol. Life Sci. CMLS 73, 3677–3691.

Ward, J.H. (1963). Hierarchical Grouping to Optimize an Objective Function. J. Am. Stat. Assoc. 58, 236.

Xu, H.P., Furman, M., Mineur, Y.S., Chen, H., King, S.L., Zenisek, D., Zhou, Z.J., Butts, D.A., Tian, N., Picciotto, M.R., et al. (2011). An instructive role for patterned spontaneous retinal activity in mouse visual map development. Neuron 70, 1115–1127.

Xu, H.-P., Burbridge, T.J., Chen, M.-G., Ge, X., Zhang, Y., Zhou, Z.J., and Crair, M.C. (2015). Spatial pattern of spontaneous retinal waves instructs retinotopic map refinement more than activity frequency. Dev. Neurobiol. 75, 621–640.

Zhang, J., Ackman, J.B., Xu, H.P., and Crair, M.C. (2012). Visual map development depends on the temporal pattern of binocular activity in mice. Nat Neurosci 15, 298–307.

