## Supplementary Figures for "Control of spontaneous activity patterns by inhibitory signaling in the developing visual cortex"

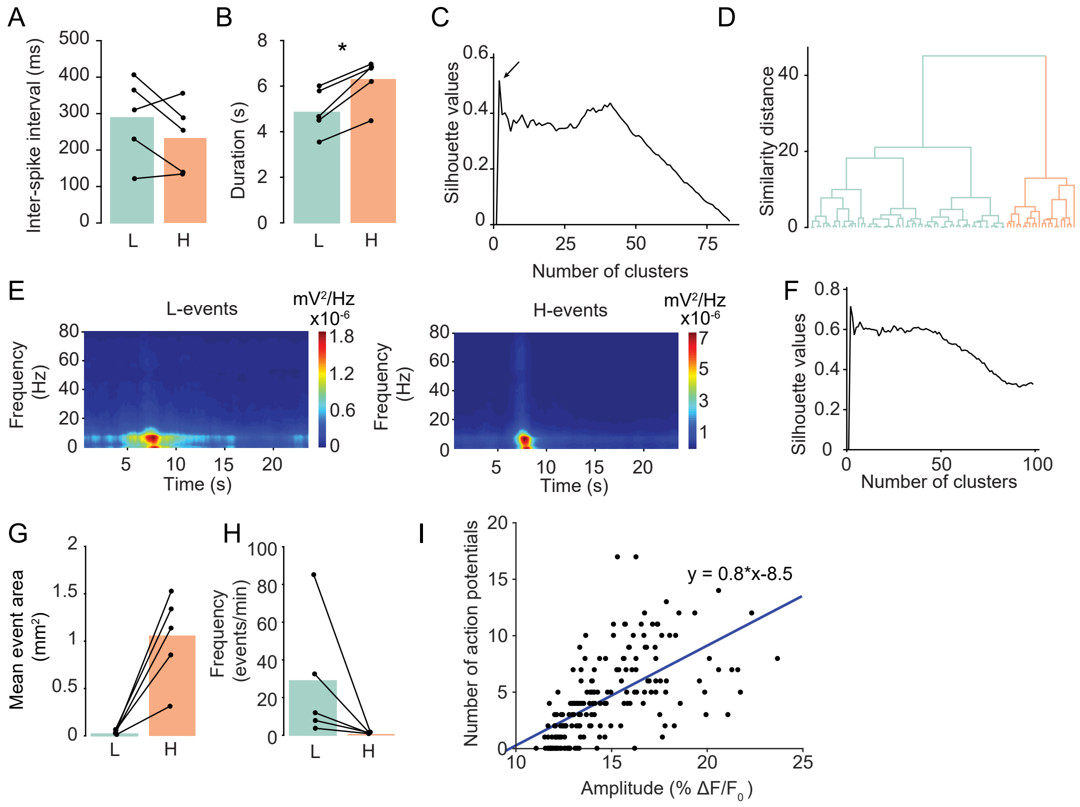


Supplementary Figure 1

A. Inter-spike-intervals within a burst are similar in L and H events (n.s., paired t-test, n = 5 animals).

B. H-events lasted significantly longer than L-events (p = 0.017, paired t-test, n = 5 animals).

C. Silhouette analysis of hierarchical clustering on the number of action potentials and the duration of depolarization shows a local peak at 2 clusters.

D. Dendrogram of hierarchical clustering performed on the duration and number of action potentials fired in an event, split into cluster 1 (L-events, blue) and cluster 2 (H-events, orange).

E. Heatmap of frequency components in the LFP. The peak frequency (L: 3.85 ± 0.2, H: 4 ± 0.12 Hz) was not significantly different (paired t-test, n = 4 animals).

F. Silhouette analysis of hierarchical clustering on the amplitude and size of events peaks at 2 clusters.

G. The event area in wide field calcium imaging is smaller during L- than during H-events.

H. The frequency of L-events is larger than H-events when imaged in wide field.

I. The relationship between the number of action potentials fired by the recorded cells and the average peak amplitude of the calcium transients in active cells in the surrounding network.

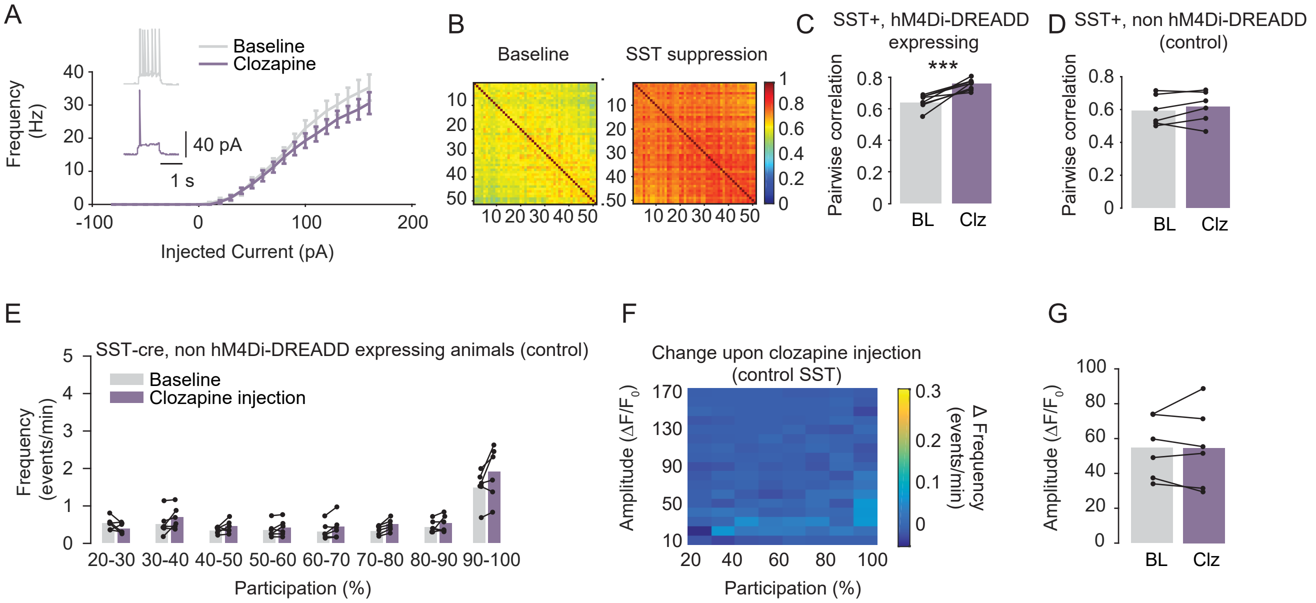


Supplementary Figure 2

A. Clozapine application in vitro significantly reduced excitability in SST+ cells expressing the hM4Di-DREADD construct (n = 8 cells, 2-ways Anova, p < 0.0001). Inset: example cell firing at 60 pA injection.

B. Spearman's rank correlation coefficient between each pair of cells in one example animal before and after clozapine administration to suppress SST neurons.

C. The mean pairwise correlation increased upon SST suppression (p = 0.0005, paired t-test, n = 8 animals).

D. No change in pairwise correlation was detected upon subcutaneous administration of clozapine in control animals not expressing the hM4Di-DREADD construct (ns, paired t-test, n = 6 animals).

E. No change in frequency was found after subcutaneous administration of clozapine in SST-cre control animals not expressing the hM4Di-DREADD construct (ns, paired t-test, n = 6 animals).

F. A heatmap showing change over various amplitudes and participation ranges shows no change after injecting the DREADD agonist clozapine in SST-cre animals not expressing the hM4Di-DREADD construct.

G. No significant change in the amplitude of events was detected before and after clozapine in SST-cre animals not expressing the hM4Di-DREADD construct.

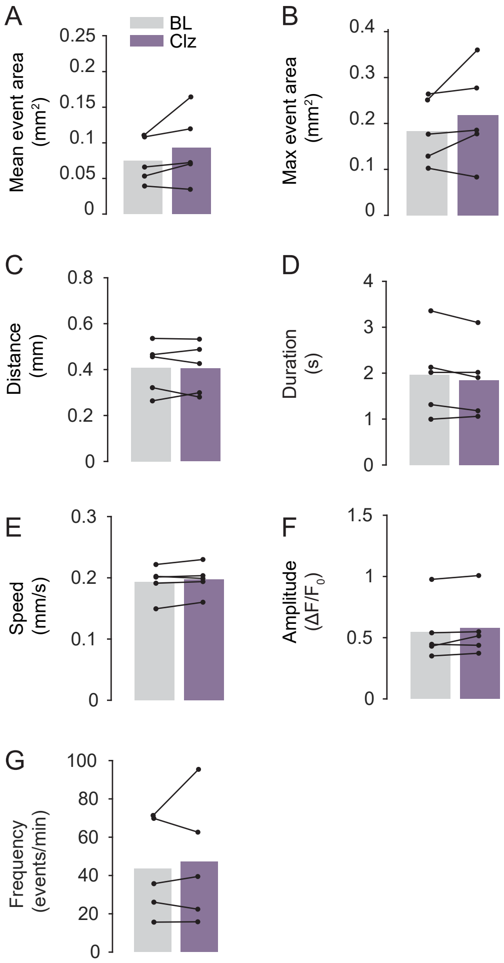


Supplementary Figure 3

A. Upon clozapine administration, the mean total area activated by an event did not change in control animals (ns, paired two-tailed t-test, n = 5 animals).

B. We detected no change in the peak size of wide-field events upon clozapine administration in control animals (ns, paired two-tailed t-test, n = 5 animals).

C. The mean distance travelled across the cortex did not change after clozapine administration in control animals (ns, paired two-tailed t-test, n = 5 animals).

D. The mean duration of events did not change after clozapine administration in control animals (ns, paired two-tailed t-test, n = 5 animals).

E. We detected no change in the speed of events upon clozapine administration in control animals (ns, paired two-tailed t-test, n = 5 animals).

F. We detected no change in the mean amplitude of wide-field events upon clozapine administration in control animals (ns, paired two-tailed t-test, n = 5 animals).

G. We detected no change in the frequency of events upon clozapine administration in control animals (ns, paired two-tailed t-test, n = 5 animals).
